# Evolutionary Action of *de novo* Missense Variants across Pathways Prioritizes Genes Linked to Autism and Predicts Patient Phenotypic Severity

**DOI:** 10.1101/158329

**Authors:** Amanda Koire, Christie Buchovecky, Panagiotis Katsonis, Young Won Kim, Stephen J. Wilson, Olivier Lichtarge

## Abstract

The pathogenicity of individual *de novo* missense mutations in autism spectrum disorder remains difficult to validate. Here we asked in 2,384 probands whether these variants exhibited collective functional impact biases across pathways. As measured with Evolutionary Action (EA) in 368 gene groupings, we found significant biases in axonogenesis, synaptic transmission, and other neurodevelopmental pathways. Strikingly, both *de novo* and inherited missense variants in prioritized genes correlated with patient IQ. This general integrative approach thus detects missense variants most likely to contribute to autism pathogenesis and is the first, to our knowledge, to link missense variant impact to autism phenotypic severity.

## Introduction

Autism spectrum disorder (ASD) is a neurodevelopmental disorder characterized by impairments in communication and social interaction, restricted and repetitive patterns of behavior (Murdoch and State 2013), and neuroanatomical abnormalities (Donovan and Basson 2016). While the most recent available data estimate that between 1/68 and 1/45 individuals are affected with autism (Christensen DL 2016), ASD is both phenotypically and genetically heterogeneous (An and Claudianos 2016). Some predictions place the number of genes involved in autism pathogenesis in the hundreds (Iossifov et al. 2012) (Betancur 2011) bordering on thousands (Liu et al. 2014) (Abrahams et al. 2013) (Yin and Schaaf 2016), and the highly multigenic nature of the disorder means that few causative genes can be identified through an excess of mutations. In the absence of any single gene responsible for the majority of ASD cases, the most commonly mutated genes only account for approximately 2% of cases each (Abrahams and Geschwind 2008) (An and Claudianos 2016). To explain additional cases, it is critical to expand analysis to interpret the collectively large number of variants in rarely mutated genes.

Although ASD has many implicated contributing factors, including environment (Durkin et al. 2008) (Cheslack-Postava et al. 2011), common polymorphisms (Gaugler et al. 2014), and inherited rare variants (Krumm et al. 2015), *de novo* variants in particular are suspected to be enriched as a class for causative mutations because they have not been subjected to generations of evolutionary selection. Analysis of *de novo* mutations in autism has largely focused on copy number variants (CNVs) (Pinto et al. 2014) (Sanders et al. 2011) (Leppa et al. 2016), single nucleotide variants (SNVs) resulting in an obvious loss of function (Wang et al. 2016) (Iossifov et al. 2014), and genes with a detectably elevated mutation rate (Sanders et al. 2012). Far less attention has been paid to the role of missense variants, whose effects on protein function are more challenging to interpret (Miosge et al. 2015) and subject to disagreement between different methods of variant impact prediction (Hicks et al. 2013). The overall role of missense variants in driving phenotype severity has also remained unclear; while strong links between mutation and lowered patient IQ have been detected for loss-of-function (LOF) and likely-gene-disrupting (LGD) *de novo* variants, as defined by the combined class of nonsense, frameshift, and splice-site mutations (Robinson et al. 2014) (Iossifov et al. 2014), studies have not yet been able to link missense mutations to the same patient presentations on a large-scale (Iossifov et al. 2014). However, individuals with ASD are more likely to carry a *de novo* missense variant than either a *de novo* loss-of-function or a *de novo* copy number variant (Iossifov et al. 2014), so the prioritization and interpretation of these variants is paramount, especially if they are revealed to be an important and understudied source of driver events.

Here, we prioritized rarely mutated, potentially causative autism genes by their *de novo* missense variants alone. Without making any *a priori* assumptions of which genes or pathways drive ASD, we tested whether groups of functionally related genes were biased toward high impact variants. To estimate the impact of each variant, we first used the Evolutionary Action (EA) equation (Katsonis and Lichtarge 2014), a state-of-the-art prediction method (Katsonis and Lichtarge 2017) that links genotype variations to fitness effects from first principles. Then to quantify mutational bias in pathways, we integrated EA over the *de novo* missense mutations of functionally related genes. This approach detected non-random mutational patterns indicative of proband-specific selection of missense variants associated with axonogenesis, synaptic transmission, and other neurodevelopmental pathways. Strikingly, in the genes prioritized by this approach, both missense *de novo* variants as well as rare inherited missense variants correlated significantly with patient IQ, demonstrating a direct relationship to patient phenotype. We concluded that pathway EA integration successfully detected the missense variants most likely to contribute to autism pathogenesis, with implications for prioritizing genes and variants and elucidating the genotype-phenotype relationship in other complex diseases.

## Results

### Characterization of the *de novo* missense variant class in ASD probands

We first assessed whether *de novo* missense variants in autism probands have, as a class, a distinct and more impactful variant profile compared to random expectation or those in matched siblings. Across 2,384 patients, we identified 1,418 missense variants affecting 1,269 unique genes and annotated the impact of the variants using Evolutionary Action (EA). Close to half of the probands (43.9%) carried a *de novo* missense variant, and the observed *de novo* missense mutation prevalence was 0.59/proband, similar to the rates reported by Neale et. al (2012) (0.58/proband) and Sanders et. al (2012) (0.55/proband). The average predicted impact of all missense variants in probands was not significantly different from what would be expected by random mutagenesis (z score = +0.13; Fig. 1A), and missense variants in probands did not have significantly higher impacts compared to their matched siblings (p = 0.23; Fig. 1B). These results suggest that the overall landscape of *de novo* missense variants in autism patients is similar to that of the matched siblings and dominated by mutations with relatively mild impact on protein fitness.

**Figure 1.**
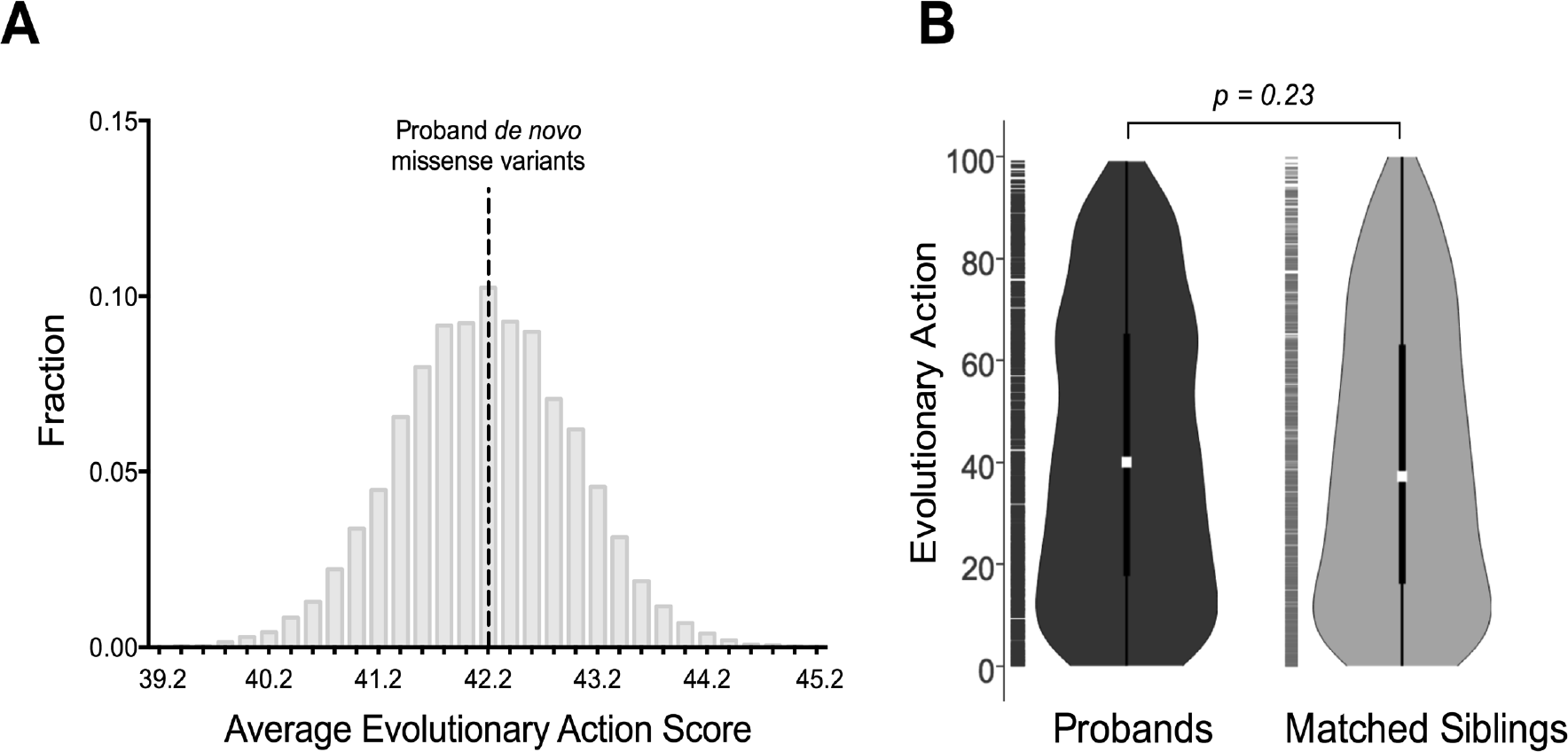
**Characterization of missense variant impact in probands and matched siblings.** (A) Proband variant impact compared to random mutagenesis. The average Evolutionary Action score of all *de novo* proband missense variants (n = 1418, in black) is superimposed upon the distribution of averages produced by 10,000 simulations of 1418 randomly selected coding missense variants. (B) Patient variant impact compared to matched siblings. The distribution of Evolutionary Action (EA) scores for missense variants in patients with matched siblings (black) versus siblings (grey) are represented by violin plots with the center dot indicating median and the center bar indicating the 25^th^ to 75^th^ percentiles of the data, and were compared statistically using a 2-sample Kolmogorov-Smirnoff test.

However, network analysis of the 1,269 genes in which missense *de novo* variants occur exposed an underlying non-random signal within this class of variants. Affected genes had significantly more protein-protein interactions in STRING (Szklarczyk et al. 2015) than would be expected by chance (p = 7.3e-12) and hundreds of GO Biological Processes were significantly enriched. Yet, the vast majority of genes under consideration exhibited these network features (Supplementary Fig. 1A) and a gene-centric interaction or enrichment approach is fundamentally limited in its ability to isolate the detected signal or stratify candidates; of the 1269 genes, 86% interact with another in the set compared to 79% expected by chance, and there is no way to identify which genes are the excess driving the significance (Supplementary Fig. 1B). For these reasons, a complementary approach to evaluating events within the missense class is necessary in order to extricate a causative subset of genes and variants.

### Prioritization of *de novo* missense variants using variant-centric pathway analysis

To pinpoint the source of the signal within the *de novo* missense class and meaningfully prioritize a subset of the missense *de novo* variants and their associated genes, we therefore pursued a variant-centric approach in which we examined patterns of variant impact across functionally related groups of genes. Genes were grouped by ontology using GO2MSIG (Powell 2014), producing 368 pathways encompassing 15,310 total genes (Supplementary Table 1), and variant impact was annotated with the Evolutionary Action (EA) method, producing impact scores on a continuous scale between 0 (minimum predicted impact) and 100 (maximum predicted impact). For the 1,792 patients with matched siblings, all 1037 patient missense *de novo* variants across 960 genes were considered. For each pathway, the EA score distribution of the *de novo* variants within the pathway was compared to the EA distribution of all other *de novo* variants. Pathways that displayed a bias toward high-impact variants and remained significant after multiple hypothesis testing were considered to be of interest, and genes that were affected by *de novo* variants and present in a significant pathway were considered prioritized genes. This approach revealed 23 significant pathways in the probands, with functions that demonstrated clear ties to nervous system development, including ‘axonogenesis’ and ‘synaptic transmission’ (Fig. 2A, Supplementary Table 2). For example, in the ‘synaptic transmission’ pathway, 49 mutations contributed from 43 individual genes produced a variant impact distribution statistically (p = 6.95e-4, q = 0.037) and visibly biased to higher EA scores (Fig. 2B). As a control, the same process was repeated using all missense *de novo* variants from the matched siblings; no pathways exhibited significant bias toward high functional impact (Fig. 2C). For subsequent analysis, genes falling into pathways with significant EA bias toward high impact mutations were grouped together into a single set of 398 ‘prioritized’ genes, and all other 562 genes with *de novo* missense variants were considered ‘deprioritized’.

**Figure 2.**
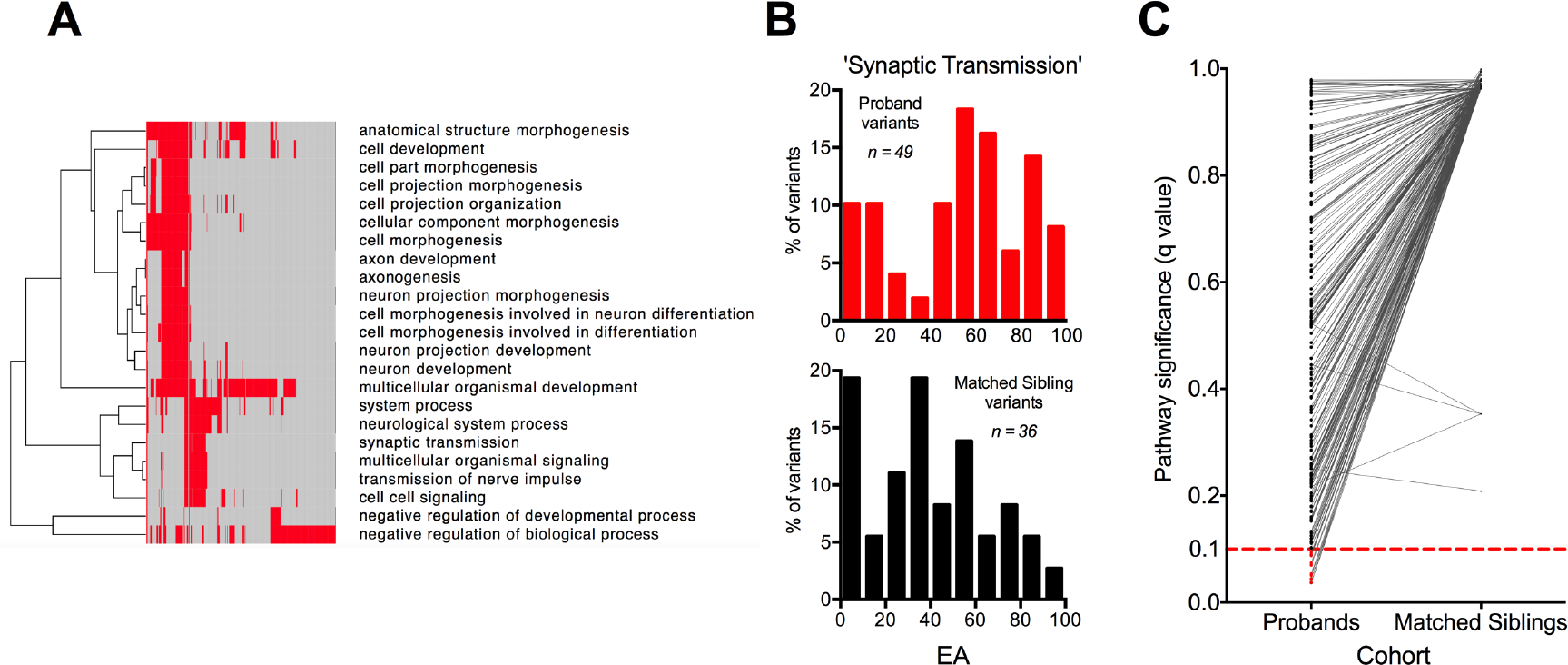
**Prioritization of *de novo* missense variants using variant impact in a pathway framework.** (A) Hierarchical clustering of significant pathways. For the 398 genes with missense mutations that were associated with at least one significant pathway, a matrix was created to denote whether the gene was (red) or was not (grey) a component of the pathway, and pathways were then grouped according to their patterns of affected genes via hierarchical clustering performed by GENE-E. (B) Evolutionary Action score distribution for ‘Synaptic Transmission’ pathway. Evolutionary Action scores for the proband variants and matched sibling variants in this pathway were binned in deciles and represented as histograms. (C) Significance of all tested pathways in patient versus matched sibling cohorts. Each point represents one of the 368 tested pathways, and is connected with a line to the same pathway in the matched cohort. The q = 0.1 significance threshold after FDR correction is represented as a dashed red line.

### EA burden of *de novo* missense variants in prioritized genes correlates with patient phenotypic severity

To determine whether prioritizing genes according to Pathway EA distributions provides a meaningful stratification between causative and non-causative genes, the variants in the prioritized genes were tested for their relationship to patient presentation, defined here by full-scale IQ. The capacity of Evolutionary Action scores alone to predict patient presentation within this prioritized gene set was tested by comparing the clinical presentations of male patients included in the initial analysis who were affected by different *de novo* missense variants in the same candidate gene. Although female probands with *de novo* missense mutations in prioritized genes contributed a minority of the data, they were highly disproportionately represented at low IQs and were analyzed separately from male patients to prevent confounding based on gender. When more than one phenotyped patient had a *de novo* missense variant in a given prioritized gene, the higher EA variant within the gene correctly predicted the patient with the lower IQ in 71.4% of paired comparisons (n = 28). Across all such cases, patients harboring the higher EA variant demonstrated significantly lower IQ overall, corresponding to an 15.2 point drop in IQ on average between the two groups (p = 0.023, paired t-test) (Fig. 3).

**Figure 3.**
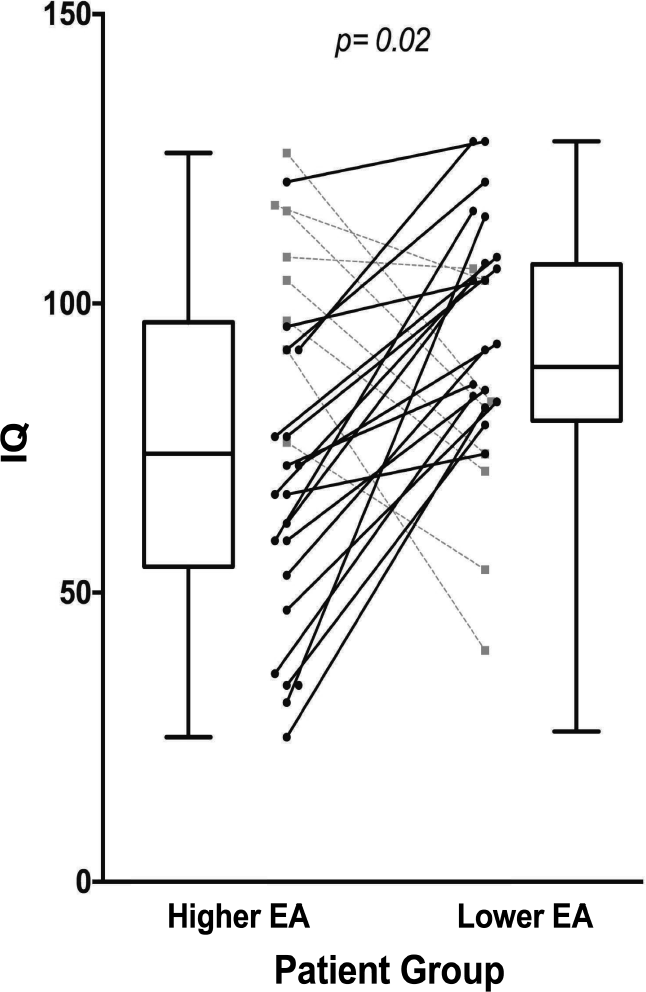
**Relationship between predicted variant impact and clinical presentation for patient pairs affected by different *de novo* missense variants in the same candidate gene.** Pairs of patients affected by different *de novo* missense variants in the same prioritized gene were identified across the 398 prioritized genes (n = 28). Within the pair, the patient with the higher variant EA score was determined, and the full-scale IQ scores of the higher-EA group were compared to the lower-EA patient group using a paired t-test. Correctly prioritized pairs are shown linked by a solid black line, while incorrectly prioritized pairs are shown linked by a dashed grey line.

To further explore the relationship between these variants and patient presentation, all male autism patients were divided into three groups corresponding to phenotypic severity: high IQ (greater than or equal to population average), low IQ (more than two standard deviations below population average, and consistent with a diagnosis of intellectual disability), and intermediate IQ. Prioritized genes were grouped together into a single set of candidate causative autism genes, and the EA score burden (sum of EA scores) of mutations in these genes was calculated for each patient and considered across the three groups. Significant differences in total variant impact were found between the three IQ groups, with the lowest IQ patient group having the highest impact mutations in the prioritized genes (p = 0.048; Kruskal-Wallis test) (Fig. 4A). This relationship between IQ and mutation EA scores was not seen when applied to all genes affected by *de novo* mutations (p = 0.58) or to genes that were not prioritized by the method (p = 0.89) (Fig. 4A).

**Figure 4.**
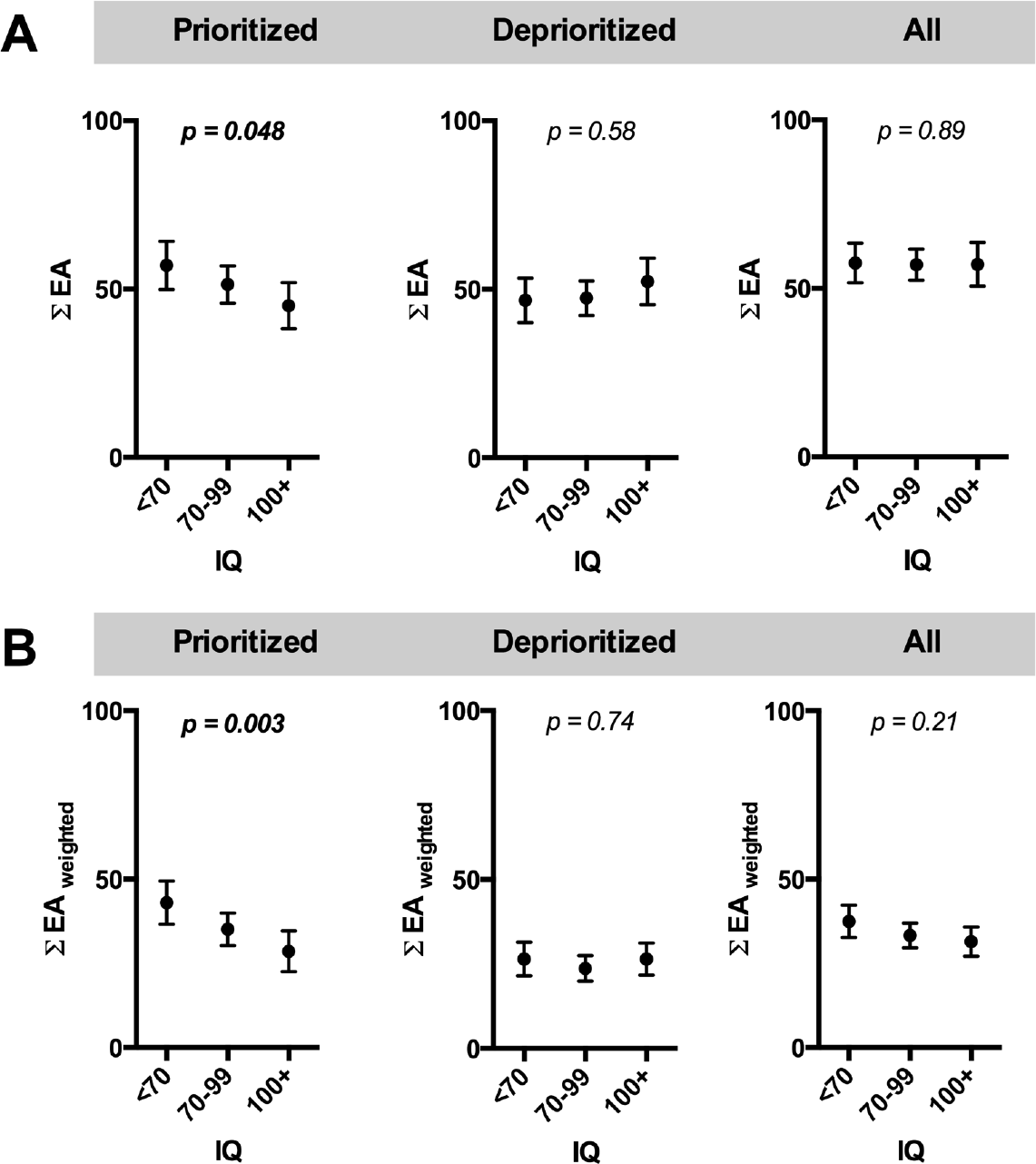
**Relationship between *de novo* EA score burden and patient IQ for prioritized and deprioritized gene groups.** Prioritized genes, deprioritized genes, and all genes with *de novo* missense variants were assessed for their relationship to patient IQ. For each male patient, the summed EA burden of *de novo* missense variants was calculated for each category, both without (A) and with (B) EA score weighting for genic intolerance to mutation. The patients were then split into three groups by their full-scale IQ score and the scores were compared using Kruskal-Wallis tests. Error bars reflect the 95% CI of the mean.

While these data show that the impact of the variant on the protein (as estimated by EA) is on its own a significant and useful predictor of patient phenotype, we next incorporated the intuitive second half of any translation between genotype and phenotype: the impact of the protein itself on human health. Although genic tolerance to mutation was not on its own predictive of phenotypic severity (Supplementary Fig. 2), adjusting the EA impact score to account for differences in genic tolerance to mutation further improved the ability of the EA score burden in prioritized genes to predict patient phenotype, and this relationship was significant both when binned (p = 0.0028; Fig. 4B) and unbinned (p = 0.013, linear regression) (Supplementary Table 3). In addition, this correlation was highly robust to the metric used to define genic tolerance to mutation (Supplementary Table 4) and was significant also when verbal or non-verbal IQ were defined as the primary outcome (Supplementary Table 5). The correlation was also robust to the impact prediction method applied, though the strongest correlations between genotype and phenotype were consistently found using EA (Supplementary Table 6). No significant relationship between patient IQ and EA score burden was found when the relevant gene set was instead considered to be all genes affected by *de novo* mutations (p = 0.21), genes that were not prioritized by the method (p = 0.74) (Fig. 4B), or genes belonging to independent gene sets of interest *a priori*, such as those enriched for expression in the brain (Uhlen et al. 2015), proposed by orthogonal methods (Darnell et al. 2011) (Parikshak et al. 2013) (Gilman et al. 2011) (Liu et al. 2014), or connected to other candidates in a protein interaction network (Supplementary Table 7).

Furthermore, while the EA score burden accounted for cases in which more than one variant of interest was detected in a patient’s exome, the results could not be explained by an uneven distribution of patients affected by multiple *de novo* variants in prioritized genes (p = 0.51; chi square test) (Supplementary Fig. 3A), and the genotype-phenotype relationship remained significant when considering only patients affected by a single variant of interest (Supplementary Fig. 3B). Female patients were assessed separately, and while their variant impact profile across prioritized genes is equally biased to high action (Supplementary Fig. 4A), the genotype-phenotype analysis is underpowered to detect a relationship of the magnitude present in male patients (Supplementary Fig. 4B) and the correlation between IQ and EA burden is not significant (p = 0.40, linear regression). These data validate that Pathway EA distributions provide a novel metric for prioritizing the set of genes most informative of clinical phenotype, and demonstrate that a clear relationship between genotype and phenotype can be detected within a subset of the *de novo* missense class.

### Prioritized gene *set* demonstrates enrichment for manually curated gold standard autism genes, uncurated literature associations, and support for novel genes

To determine whether prioritization using Pathway EA distributions captures established knowledge, we next compared our prioritized gene set to SFARI’s list of manually curated gold standard genes for autism. We considered SFARI categories 1-3 (‘high confidence’, ‘strong candidate’, and ‘suggestive evidence’) to be a gold standard for comparison. We then quantified the overlap of our gene lists to SFARI and found that the prioritized genes were highly enriched for genes in the ‘gold standard’ SFARI gene set compared to deprioritized genes (35/398 vs. 14/562; p < 0.0001, Fisher’s exact test). These data show that prioritization using Pathway EA distributions preferentially captures curated knowledge. To distinguish whether novel genes were contributing to the relationship between genotype and phenotype, we next tested the ability of EA burden in prioritized genes to predict patient phenotype when the gene was either supported by the high-confidence curated gold standard or unscored by SFARI (‘novel’). For each subset of the prioritized genes, patients with *de novo* variants in these genes were split into two groups based on whether their burden was above or below the mean of all such patients. Across all prioritized genes, the patient group with above-average EA burden demonstrated significantly lower IQ scores corresponding to an ∼8 point drop in IQ (p = 0.006) (Fig. 5). The difference became more pronounced when restricting to prioritized genes also in the ‘gold standard’ SFARI gene set, with average IQ a full 30 points lower in the patient group with higher EA burdens (90.3 vs 60.3, p = 0.0015), 5 points more than would be found when considering SFARI ‘gold standard’ genes without the aid of prioritization (Supplementary Fig. 5). However, the majority (84.5%) of prioritized genes were not placed into any category by SFARI curation, and a significant ∼6.5 IQ point difference between the groups persisted when considering only these unannotated genes (p = 0.03).

**Figure 5.**
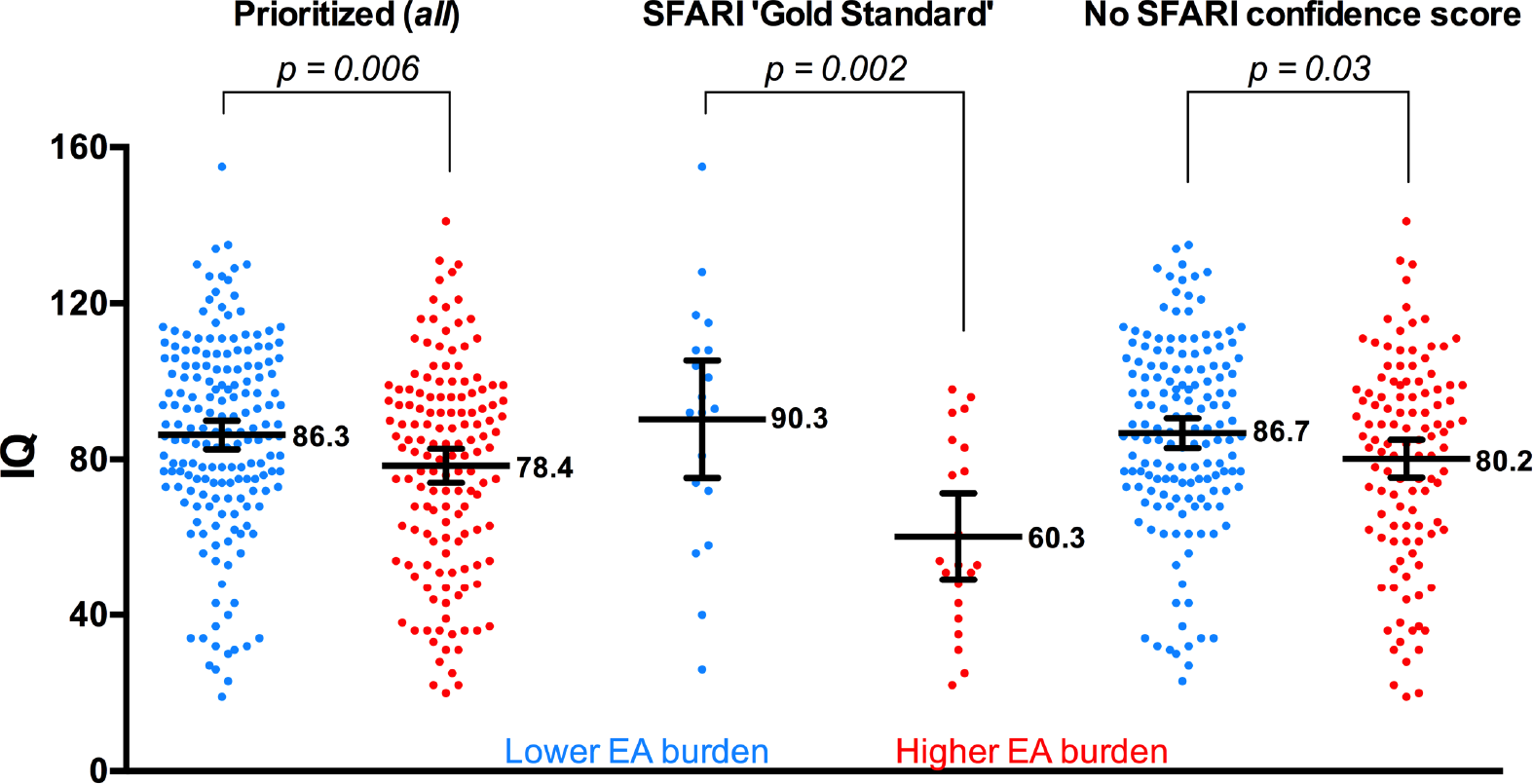
**Effect of SFARI curation confidence and prioritization status on the relationship between genotype and phenotype.** For each of three gene sets of interest (All prioritized genes, prioritized genes overlapping with a SFARI ‘Gold Standard’ designation (SFARI categories 1-3), and prioritized genes without a SFARI confidence score), the gene set EA burdens of all male patients with at least one *de novo* missense variant within the gene set were averaged, and patients were split into ‘Higher EA burden’ and ‘Lower EA burden’ groups based on whether their score was above or below the average burden, respectively. Groups were compared statistically with an unpaired t-test, and the mean and 95% CI interval of the mean for each group is displayed overlaying all IQ scores for patients in the group.

We next re-performed the analyses with a more stringent definition of novelty, comparing our prioritized gene set to an uncurated assessment of the current published literature. We defined genes with at least one association in Pubmed between the gene name and the term ‘autism’ as being supported by the literature, and found that prioritized genes were significantly enriched for literature support compared to deprioritized genes (p < 0.0001, Fig. 6A). Amongst all genes with literature support, those that were prioritized had a larger number of associations per gene (p = 0.007, Fig. 6B), indicating more extensive support. Moreover, while prioritized genes with literature support exhibited a significant relationship between IQ and EA burden (Fig. 6C), the deprioritized genes with literature support did not (Fig. 6D), suggesting that associations with deprioritized genes may be false positives. We then tested the ability of EA burden in prioritized genes to predict patient phenotype when the gene was either supported or unsupported by the literature. When considering prioritized genes with literature support, patients with an above-average EA burden had IQ scores ∼11 points lower than those with below-average EA burdens (p = 0.01, Fig. 6E); when instead considering prioritized genes with no literature associations to autism, the same trend was seen with a significant decrease in IQ of over 7 points (p = 0.04, Fig. 6E). These data support that the novel prioritized genes contribute to the significant relationship detected between genotype and patient phenotype, even when the threshold for determining novelty is made highly stringent and reflective of current scientific knowledge. Therefore, the novel set prioritized by Pathway EA distributions is likely to contain a number of genes that are genuinely causative and informative of patient phenotype.

**Figure 6.**
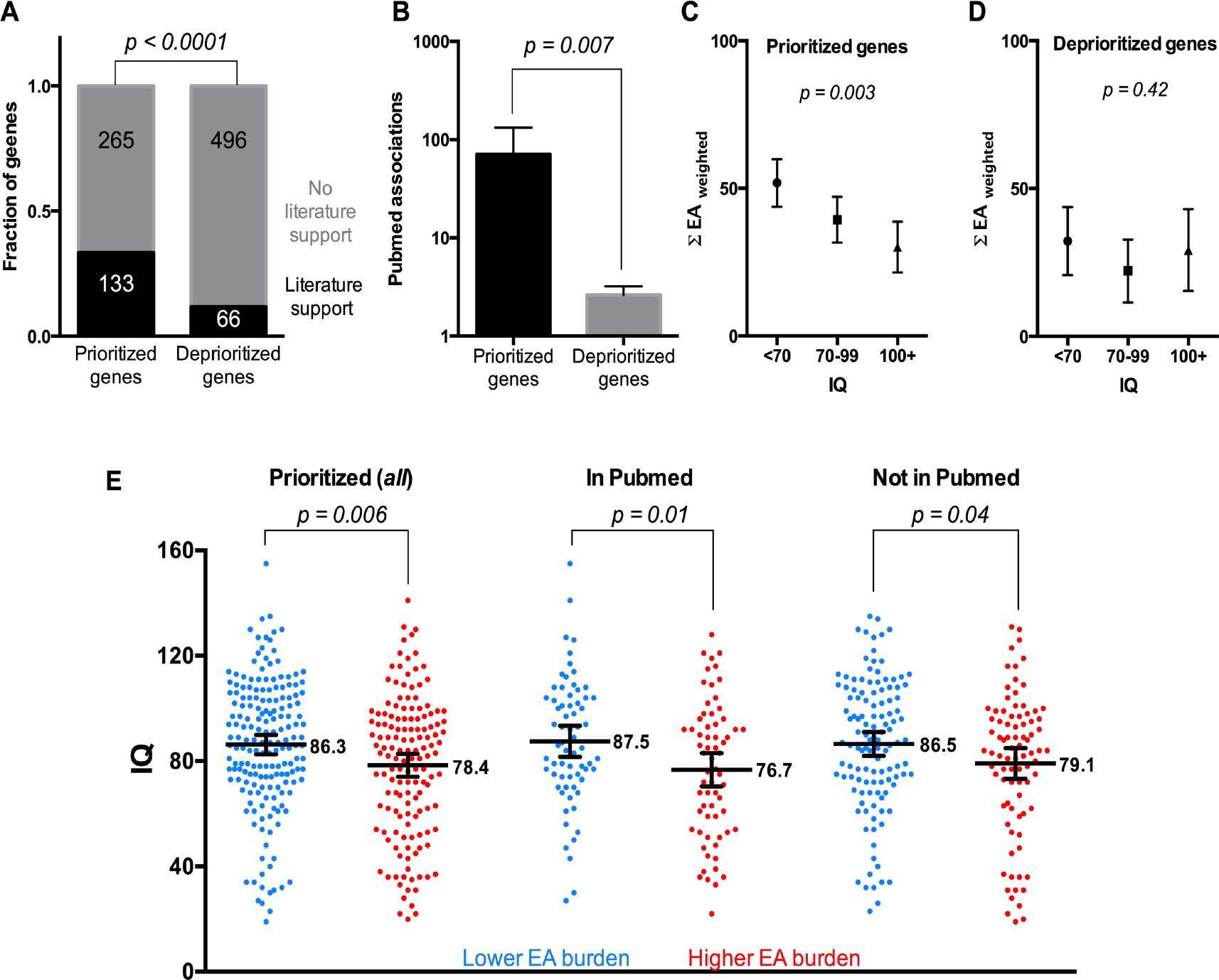
**Uncurated literature associations to autism and the effect of literature and prioritization status on the relationship between genotype and phenotype.** (A) Enrichment of prioritized gene set for associations with autism in Pubmed compared to deprioritized gene set. (B) All genes with *de novo* variants and support in the literature were separated by prioritization status and the numbers of Pubmed associations to autism for the genes in each category were compared with an unpaired t-test. (C) *de novo* missense variants in prioritized genes with literature support (n = 133) were assessed for their relationship to patient IQ. Male patients were then split into three groups by their full-scale IQ score and their weighted EA burdens were compared using Kruskal-Wallis tests. Error bars reflect the 95% CI of the mean. (D) *de novo* missense variants in deprioritized genes with literature support (n = 66) were assessed for their relationship to patient IQ as above. (E) For each of three gene sets of interest (All prioritized genes, prioritized genes with at least one Pubmed association to autism, and prioritized genes without a Pubmed association to autism), the gene set EA burdens of all male patients with at least one *de novo* missense variant within the gene set were averaged, and patients were split into ‘Higher EA burden’ and ‘Lower EA burden’ groups based on whether their score was above or below the average burden, respectively. Groups were compared statistically with an unpaired t-test, and the mean and 95% CI interval of the mean for each group is displayed overlaying all IQ scores for patients in the group.

### EA score burden of rare and low-frequency inherited variants in prioritized genes also correlates to phenotype severity

Given that the impact of *de novo* mutations in the candidate causative genes correlated to patient presentation, we next tested whether rare inherited variations in these same genes exhibited a similar relationship with IQ. We considered rare and low-frequency inherited variants (MAF<0.05) that were detected in at least one parent, but were not inherited by the healthy sibling. For each patient, we calculated the inherited EA burden in the candidate genes as the summation of all EA scores in these variants after adjustment for gene-specific tolerance to mutation. We found there was a significant correlation between IQ and inherited variant EA burden as well, with high IQ patients having a lower inherited EA burden in the prioritized gene set (p = 0.0005; Fig. 7A), while there was no relationship between EA burden and IQ when considering genes that were not prioritized (p = 0.26), or that were low-confidence SFARI genes (SFARI categories 4-6; p = 0.83); the same relationships could be found when limiting the MAF cutoff to more stringent definitions of rare variant status (Fig. 7B). Incorporation of the *de novo* variants into the EA burden increased significance further (p = 0.0003). These data show that within the prioritized gene set, rare inherited variants also link genotype to phenotype and are likely contribute to clinical severity in addition to the *de novo* variants in these genes.

**Figure 7.**
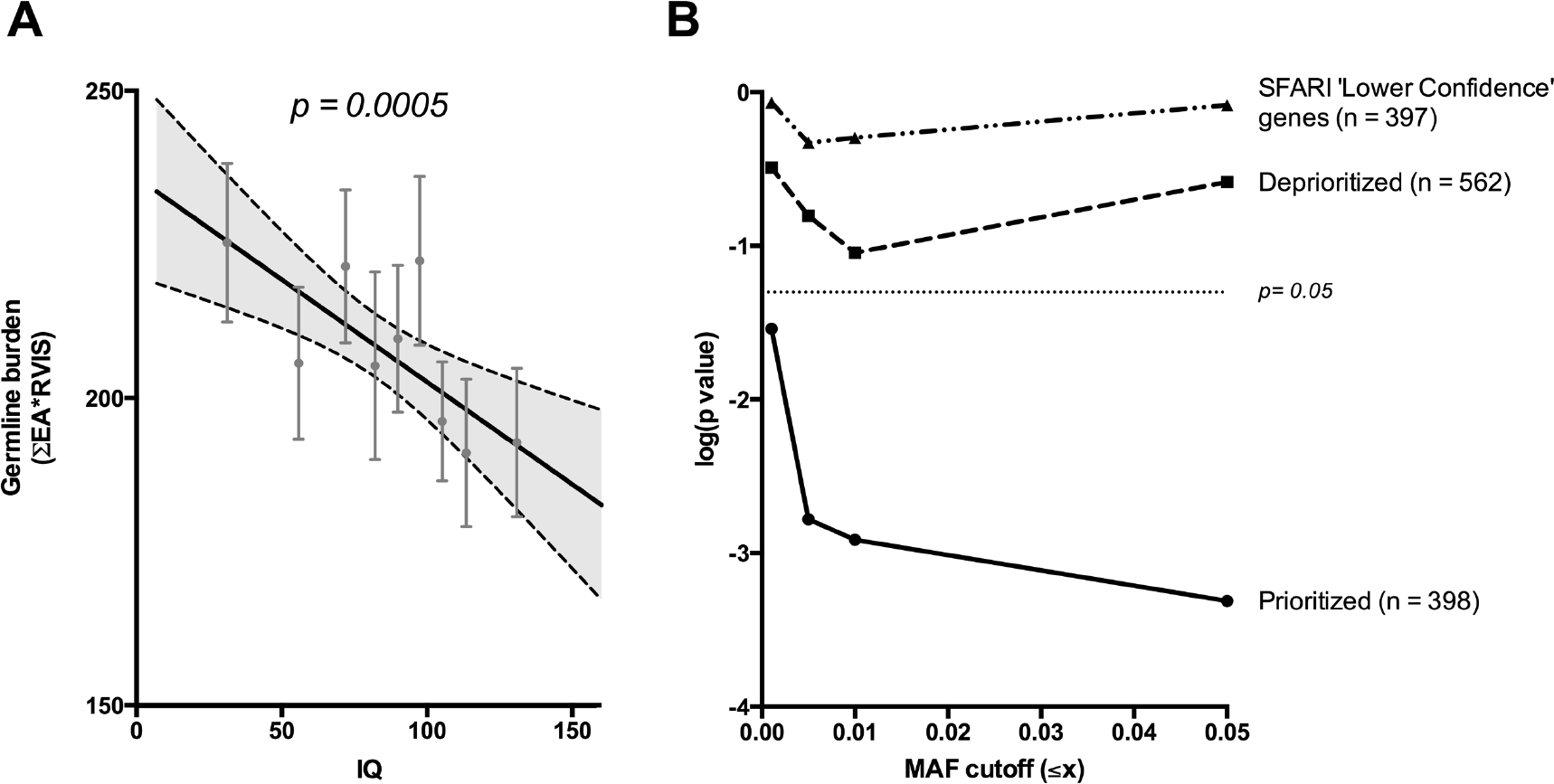
**Relationship between inherited EA score burden and patient IQ for prioritized and deprioritized gene groups** (A) For each male patient, rare and low-frequency inherited variants (MAF < 0.05) that were detected in at least one parent, but not inherited by the healthy sibling, were identified across prioritized genes and the inherited EA burden was calculated as the summation of all EA scores of these variants after weighting for genic tolerance to mutation. The line indicates the linear regression across all points and the shaded grey area represents the 95% CI; the p-value displayed corresponds to the significance of the regression. For visualization purposes, the patients were also sorted by IQ and divided into nine equal groups; the average burden and IQ of each group is overlaid upon the regression and error bars indicate the standard error of the mean. (B) Log(p-values) of the linear regression of IQ and inherited EA burden as the MAF threshold is increasingly restricted to lower frequencies, for prioritized genes, deprioritized genes, and lower-confidence SFARI genes.

## Discussion

Our data show for the first time that *de novo* missense impact signatures can be used to elucidate causative pathways in a complex multigenic disease and prioritize variants that stratify disease severity. Here, using sequencing data from 2,384 individuals diagnosed with autism, we hypothesized that affected cohort-specific selection for large variant fitness effects within a group of functionally related genes implied an association of those pathways and genes to ASD. We observed significant impact signatures in 23 pathways, including axonogenesis, neuron development, and synaptic transmission, among others. The mutated genes from these pathways were enriched for literature associations to autism and are highly consistent with pathways of importance derived from analyses of CNV and LOF variant data, (Gilman et al. 2011) (Pinto et al. 2014) (Glessner et al. 2009) suggesting that the putative causative missense SNVs identified in this study operate through mechanisms similar, rather than orthologous, to well-documented processes involved in autism etiology. Excitingly, novel genes from prioritized pathways comprise the majority and, like known genes, exhibit a genotype-phenotype relationship informative of patient presentation.

To the best of our knowledge, this study is the first to directly link missense variant impact to autism phenotypic severity. Although IQ cannot reflect all possible aspects of patient phenotypic severity, it correlates strongly to behavior-based observer-rating scales that encompass diverse areas of autistic symptomatology (Nishiyama et al. 2009), and repetitive behaviors in patients (Richler et al. 2007), therefore providing a relevant index of ASD severity. Past work relating *de novo* variants to IQ across ASD patients has focused almost exclusively on CNV and loss-of-function (LOF) variants, with studies finding a significant relationship between IQ and the *de novo* mutation rate of likely gene disrupting (LGD) variants (Robinson et al. 2014) (Iossifov et al. 2014) as well as CNVs and truncating SNVs (O’Roak et al. 2012). However, when these same studies assessed missense variants, no correlation with intellectual disability was found, even after restricting to recurrent missense variants (Iossifov et al. 2014) (O’Roak et al. 2012). While our initial assessments of the *de novo* missense class agreed with others who have reported that the overall impact of *de novo* missense variants in ASD does not differ substantially from expectations (Neale et al. 2012) (Sanders et al. 2012), we found that this collective profile did not preclude the detection of gene and variant subsets with mutational signatures indicating a significant genotype-phenotype relationship. We observed a 30 point IQ decrease in patients with above-average missense impact burdens across the highest confidence candidates, and a 7.4 point IQ decrease in patients with above-average missense impact burdens across the most novel candidates. Additionally, we demonstrated that different variants within the same candidate gene can be linked to phenotypic outcomes through their predicted EA impact on protein fitness. Furthermore, a modest but highly significant correlation between rare inherited missense burden and IQ when considering the prioritized genes indicates that these genes may contribute to autism etiology through avenues beyond *de novo* variation.

On the whole our results suggest that *de novo* missense variants, especially those with high impact affecting important genes in neurologic pathways, have the potential to influence patient presentation even if they or the genes in which they occur have not been previously linked to autism in the literature; however, lower-impact missense variants in a gene should not be assumed to produce a similar effect even if the gene or pathway has been previously associated with autism. These findings have implications for clinical interpretation of *de novo* missense variants of unknown significance in patients diagnosed with autism, which in turn can improve estimations of recurrence risk in siblings by helping to clarify whether a patient’s *de novo* missense variant influences their presentation or is merely incidental. In the future, larger cohorts and additional sequenced trios will allow for refinement of the observed genotype-phenotype relationship into a clinically valuable outcome predictor, and clarify whether missense variants in female patients with autism demonstrate the same relationship to clinical presentation.

In addition, our results have implications for laboratory testing by suggesting which genes and variants to prioritize for experimental validation and inclusion into the SFARI gold standard. One gene with a single missense variant in the cohort, CAMK2A, was not included in SFARI Gene when this study was completed and had minimal literature support for an association to autism but was prioritized by the pathway-EA integration as part of the synaptic transmission pathway. The detected variant in this gene has very recently been shown to decrease excitatory synaptic transmission in cultured neurons and produce aberrant behavior including social deficits and increased repetitive behavior in mice with a knock-in of the variant (Stephenson et al. 2017), and has since been incorporated into SFARI Gene. Though at the moment a single example, pathway-EA can prospectively aid ongoing large-scale experimental efforts to test the functional effect of *de novo* missenses mutation detected in major trio studies.

In a broader scope, the discovery of a genotype-phenotype relationship through the integration of mutation impact and gene importance scores is an approach whose success has implications for evolutionary theory at large. The mathematical underpinning behind using EA distributions to identify pathways and genes of interest is founded on the assumption of an evolutionary fitness function that maps genotypes to phenotypes in the fitness landscape, but which is not directly calculable. Differentiation of this fitness function yields the EA equation to predict variant impact, in which the perturbation of the fitness landscape is equal to the product of the evolutionary fitness gradient, estimated by Evolutionary Trace (Lichtarge et al. 1996), and the substitution log-odds of the amino acid change(Katsonis and Lichtarge 2014). These values are calculable from sequence data and predictions have been shown to correlate well to experimental assessments of protein fitness (Gallion et al. 2017) (Katsonis and Lichtarge 2014), consistently outperform machine learning methods (Katsonis and Lichtarge 2017), and to stratify patient morbidity (Katsonis and Lichtarge 2014) and mortality (Neskey et al. 2015) in other disease contexts. Here this EA theory is extended by considering the distribution of variant EA scores over a pathway. Such distributions amount, in effect, to integrating the EA equation across the pathway to recover the original genotype-phenotype relationship. Significant distributions indicate a nonrandom genotype-phenotype relationships. As we show here, this new evolutionary calculus in fitness landscapes can, in practice, identify candidate phenotypic driver genes and the relationship between variant impact and patient clinical outcome. Though it is applied here to the ASD phenotype, the pathway EA approach is highly generalizable to other multigenic diseases and phenotypes and can be applied to germline and *de novo* mutations alike.

## Methods

### Data acquisition

Variant call files (.vcfs) produced by the Simons Simplex Collection (SSC) were downloaded from NDAR (Study 349); this exome data encompassed 2,392 families and used FreeBayes SNV calling performed by Krumm et al. at the University of Washington. Phenotype data for the associated patients were obtained from the same source.

### De novo variant calling and quality assessment

Variants were called as *de novo* if the proband call was heterozygous with a depth higher than 10, alternate allele fraction of 0.3 or higher, and average alternate allele quality of 15 or higher; the same position was required in both parents to have a depth of at least 30, at least 95% of reads supporting a reference call, and no more than 5 reads supporting a non-reference call. These thresholds produced a set of *de novo* variants indicating high quality (Ti/Tv=2.64) and an absence of negative selection (lambda=0.009) (Koire et al. 2016). Using this procedure we identified *de novo* variants in both patients and siblings. Eight families were excluded from downstream analysis due to an excessive number of apparent de novo sequence events in either the patient or sibling, suggesting an apparent sample swap or a non-biological relationship between the children and at least one parent. In order to focus on genes that are infrequently mutated we did not consider genes with more than three missense mutations, which notably included well-documented autism driver SCN2A, and analyzed only non-recurrent variants.

### Network/Gene Set Enrichment analysis of genes affected by *de novo* variants in patients

Protein-protein interactions were defined by the *Homo sapiens* STRING v.10.0 network (Szklarczyk et al. 2015) using the aggregate score of all evidence types and were considered as interactions if they had ‘medium confidence’ or higher (interaction score ≥ 0.4). Enrichment tests for protein-protein interactions, as well as gene set enrichment analysis for GO Biological Processes, were performed through the STRING graphical user interface. Gene sets were considered significantly enriched at the default q<0.05 threshold reported by STRING.

### Annotation of missense variants with Evolutionary Action

The impact of missense variants on protein fitness was computed with the Evolutionary Action (EA) equation, which has won multiple CAGI challenges in 2015, 2013, and 2011 (Cai et al. 2017). Briefly, this equation follows from viewing evolution as a differentiable mapping, *f* of genotypes (*γ*) onto the fitness landscape (*φ*), so that:

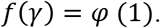

Differentiation then leads to the EA equation:

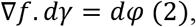

where ∇*f* is the evolutionary gradient in the fitness landscape, *dγ* is a genotype perturbation such as a mutation, and *dφ* is the fitness effect. In practice, (2) is approximated to first order. For a substitution from amino acid type *X* to type *Y* at a protein residue, *r_i_*, the evolutionary gradient ∇*f* reduces to 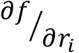, the mutational sensitivity at *r_i_* also equivalent to its evolutionary importance defined by the Evolutionary Trace method (Lichtarge et al. 1996) (Mihalek et al. 2004). To estimate *dγ*, we use odds of amino acid substitution from *X* to *Y*. This approach produced scores on a continuous scale between 0 and 100, where a higher value indicated a larger predicted impact on protein fitness resulting from the amino acid substitution. When a variant affected multiple isoforms of a protein, the impact score was averaged across all affected isoforms. Evolutionary Action calculations are described at greater length in the original publication of the method (Katsonis and Lichtarge 2014).

### Identification of gene groups with bias toward impactful missense variants

Gene groups were defined using Gene Ontology (GO) terms customized by GO2MSIG (Powell 2014); customization was specific to *Homo sapiens* and ensured at least 500 genes in each group. This approach produced 368 pathways encompassing 15,310 total genes (Supplementary Table 1). Gene groups with a collective variant bias toward high impact were identified by examining the EA score distributions of their missense *de novo* variants. For each pathway, the EA score distribution of the *de novo* variants within the pathway was compared to the EA distribution of all other *de novo* variants using a one-sided Kolmogorov-Smirnov test. Groups which were significant after FDR with q<0.1 were considered significant (Supplementary Table 2). This analysis was performed using missense *de novo* variants from 1,792 patients with matched siblings, and then repeated using missense *de novo* variants from the 1,792 matched siblings.

### Relating Evolutionary Action scores in prioritized genes to patient phenotype

Although female probands with *de novo* missense mutations in prioritized genes contributed a very small fraction of the data (<1/7), they were highly disproportionately represented at low full-scale IQ scores (42% with IQ < 70 vs. 26% for male probands) and were analyzed separately from male patients to prevent confounding based on gender. Autism patients were divided into three groups by phenotype severity as defined by high full-scale IQ (greater than or equal to population average), low full-scale IQ (more than two standard deviations below population average, and consistent with a diagnosis of intellectual disability), and intermediate full-scale IQ. Genes falling into pathways with significant bias toward high impact mutations were grouped together into a single set of prioritized candidate autism genes, and we considered the EA scores of mutations in these genes across the three groups for all binned analyses.

For each patient the sum of the Evolutionary Action scores of *de novo* variants in their affected candidate genes was calculated. We used the Residual Variation Intolerance Score (RVIS) (Petrovski et al. 2013) as our main measure of genic sensitivity to mutation; RVIS scores were converted with the equation *mutation intolerance score = ((100-RVIS%)/100)* in order to lie on a scale from 0-1 with 1 indicating maximum intolerance to mutation. For each gene, variant EA scores were then adjusted by the intolerance score such that *adjusted EA= EA*mutation intolerance score*, and the total patient burden was recalculated with adjusted EA scores in place of the raw EA scores. For comparison, we also substituted raw ExAC LOF Constraint Metric (pLI) and ExAC Missense Constraint Metric scores as alternative measures of genic intolerance to mutation (Lek et al. 2016) (Supplementary Table 4), nonverbal and verbal IQ scores as alternate measures of phenotypic severity (Supplementary Table 5), other prioritization approaches (Supplementary Table 7), and CADD (Kircher et al. 2014), SIFT (Ng and Henikoff 2003), and Polyphen-2 (Adzhubei et al. 2010) scores as alternate measures of variant severity (Supplementary Table 6).

For analysis of inherited germline variants, we considered low-frequency inherited variants (MAF<0.05) in prioritized genes that were observed in at least one parent, but were not inherited by the healthy sibling (using the same thresholds to confirm absence of the variant as were used for parent calls when determining *de novo* variants). The inherited EA burden for each male patient was calculated as the summation of all EA scores of these variants after weighting for genic tolerance to mutation. The minor allele frequencies of variants were obtained from ExAC (Lek et al. 2016).

### Comparison of prioritized genes to published knowledge

Genes with at least one association in Pubmed between the gene name and the term ‘autism’ were defined as being supported by the literature, while genes with no search results returned were defined as lacking literature support. These values were obtained automatically using a Biopython script on 10/4/2016. SFARI gene annotations were obtained from SFARI Gene and the SFARI Gene Scoring Module (Abrahams et al. 2013) on 1/13/2017.

## Data Access

Variant call files (.vcfs) were downloaded from NDAR (Study #349); approved researchers can also obtain the underlying SSC population dataset described in this study (https://sfari.org/resources/autism-cohorts/simons-simplex-collection) by applying at https://base.sfari.org

## Acknowledgments

This work was supported by the National Institutes of Health [grant numbers GM079656-8 to O.L., DE025181 to O.L., GM066099 to O.L.]; the National Science Foundation [grant number DBI1356569 to O.L.], and the Defense Advance Research Project Agency [grant number N66001-15-C-4042 to O.L]. A.K. is supported by RP160283 - Baylor College of Medicine Comprehensive Cancer Training Program, and the Baylor Research Advocates for Student Scientists (BRASS).

We are grateful to all of the families at the participating Simons Simplex Collection (SSC) sites, as well as the principal investigators (A. Beaudet, R. Bernier, J. Constantino, E. Cook, E. Fombonne, D. Geschwind, R. Goin-Kochel, E. Hanson, D. Grice, A. Klin, D. Ledbetter, C. Lord, C. Martin, D. Martin, R. Maxim, J. Miles, O. Ousley, K. Pelphrey, B. Peterson, J. Piggot, C. Saulnier, M. State, W. Stone, J. Sutcliffe, C. Walsh, Z. Warren, E. Wijsman), and appreciate obtaining access to phenotypic data on SFARI Base.

